# Ionic environment and nutrients affect the *cysB*-dependent conditional susceptibility of *E. coli* to mecillinam

**DOI:** 10.1101/2025.09.27.678940

**Authors:** L Mancini, M Kals, J Kotar, P Cicuta

## Abstract

Antibiotic efficacy is influenced by environmental context. Urinary *E. coli* isolates lacking a functional *cysB* gene have been shown to be resistant to the *β*-lactam antibiotic mecillinam under high-osmolarity conditions, suggesting that increased fluid intake might synergize with drug treatment. In this study, we further explore *E. coli* susceptibility to mecillinam across growth conditions. We show that while osmolarity has a mild protective effect on *cysB* knockout and reference strain, it is the presence of certain salts that leads to the resistant phenotype in the mutant. Such conditional resistance is specific to mecillinam and rapidly wanes when salts are removed. In agreement with our previous work showing that slow growing cells escape mecillinam-induced bursting by maintaining a small size, for both strains we observe low susceptibility to mecillinam in scarcely nutritious pooled human urine. As we supplement urine with nutrients, we observe a sharp transition to susceptibility when growth rate surpasses 0.6 h^*−*1^. Growth in the presence of salt or uncharged osmolites decreases cell size of both reference strain and *cysB* knockout offering a potential explanation to the observed mild protective effects of osmolarity. Our findings give insights into the mechanism of *cysB* dependent susceptibility to mecillinam and, due to the dual impact of nutrients and osmolites on treatment, allow us to reformulate ideas on potential diet recommendations. While increased fluid intake will decrease urine osmolarity, it will also reduce nutrient content, limiting mecillinam efficacy. We suggest instead that reducing sodium intake during mecillinam treatment might avoid this trade-off.

## INTRODUCTION

Urinary tract infections affect more than 400M people per year worldwide (1) with high recurrence (2). They are often treated with *β*-lactams, a class of cell wall-targeting antibiotics whose efficacy depends on the cells’ growth rate and on the osmotic properties of their environment. Bacterial cell walls grow as a result of the coordination between synthesis and hydrolysis (3; 4). *β*-lactams bind to the penicillin binding proteins (PBPs) that produce and crosslink the cell wall, leading to an imbalance in favour of hydrolytic processes that eventually results in compromised walls. Because these processes are much more active during periods of active growth, the drugs have scarce efficacy when growth is arrested, resulting in a strong antagonism with bacteriostatic antibiotics (5; 6) and poor efficacy against dormant cells (7). The cell wall is a major load-bearing structure in bacterial cells (8) and, further to protecting it from mechanical stresses from the outside, it supports its turgor pressure. Treatment with *β*-lactams can lead to the formation of holes in the cell wall, through which the cytoplasm can bulge, swell and rupture (9). The stability of cells with compromised walls depends on osmolarity (10; 11). Cells that have lost their cell wall remain vital in media with high osmolarities, in the form of L-shapes or spheroplasts (12). The main mechanism of resistance to *β*-lactams is the enzyme betalactamase, which inactivates the drugs by hydrolisis (13).

Among *β*-lactams, mecillinam is one of the least susceptible to betalactamase activity (14) and it is used to treat urinary tract infections. As well as for its stability, mecillinam is special in its class because rather than having high affinity for a number of penicillin binding proteins (PBPs), it preferentially binds to the enzyme PBP2 which catalyzes the cross-linking of peptidoglycan strands within the cell wall (15). Cell death due to mecillinam occurs as a consequence of progressive cell swelling (SI Fig. 1) and we have shown that poor media or mechanical constriction lead to cell survival by limiting swelling (16). In clinical settings, despite the reduced susceptibility to betalactamases, resistance still emerges and most resistant isolates carry mutations that inactivate *cysB*, a global regulator of cysteine biosynthesis (17). The mechanism of resistance remains unknown, but *cysB*-dependent susceptibility in urine has been found to be linked to urine concentration (18), with resistance only appearing at high osmolarity. The bactericidal action of *β*-lactams is known to be linked to the osmolarity of the cell and of its milieu, because these determine the likelihood of bursting of the cytoplasmic blebs that develop from cell wall damage (9).

In this work, we further characterize the relationship between environmental conditions and mecillinam resistance in the Δ*cysB* background. Our results reveal that while increased osmolarity has a certain protective effect on *E. coli* against mecillinam, it is only certain salts that enable high-level resistance, pointing to a distinct mechanism. We show that this mechanism is unique to mecillinam, as Δ*cysB* cells are susceptible to other *β*-lactams and resistant to novobiocin (19) independently of salt. We further show that incubating cells or the drug with salt before treatment is administered does not influence efficacy, indicating a mechanism with relatively rapid dynamics. Lastly, as we observe that slow growing cells, including those grown in native urine, are scarcely susceptible to mecillinam, these findings support our recently-proposed model of geometry-dependent mecillinam killing efficacy.

## METHODS

### Bacterial strains and growth conditions

Experiments were carried out using *E. coli* BW25113 and its Δ*cysB* derivative, which was retrieved from the Kejo collection (20). Before experiments, freshly streaked single colonies were grown overnight in LB (0.5% Yeast Extract, 1% Bacto tryptone, 0.05% NaCl) in flasks incubated at 37^°^C with shaking at 200 rpm. LB solutions were prepared in all cases starting from LB_*L*_ (5 g/l yeast extract, 10 g/l tryptone, 0.5 g/l NaCl). To this, osmolites were added such that the final osmolarity matched to that of 2.5, 5 and 10 g/l total NaCl (SI table 1). M63 (13.6 g/L KH_2_PO_4_, 0.5 mg/L FeSO_4_ · 7 H_2_O, 0.5 g/L MgSO_4_ · 7 H_2_O, 1.27 mg/L thiamine, 2.64 g/L (NH_4_)_2_SO_4_, and 0.5% w/v glucose, final pH: 7.2) was prepared as before (21), while MOPS buffer, EZ amino acid and vitamin mixes, and AGCU nucleobases solutions were purchased from Teknova, US. Aliquots of these commercial components were thawed just before each experiment and added directly to M63 and pooled human urine (Stratech, UK). The final concentration of M63 in M63+AGCU was 90%, in M63+EZ was 80% and in M63+EZ+AGCU was 70%. Urine was used at a concentration of 50% in all experiments except those in Fig. 4A in which it was undiluted. Mecillinam stocks were freshly prepared in water.

### Optical density measurements

Cells from overnight cultures were inoculated in either 96 or 384-well flat-bottom plates (Costar, UK) filled with 200 or 50 *µ*l of medium, respectively, at a 1:100 dilution. Optical density measurements over time were carried out in a Spectrostar Omega microplate reader (BMG, Germany) at 37^°^C, with 700 rpm shaking (double orbital mode), with the lid on.

### Brightfield microscopy

Imaging was performed as described before (22; 23). Briefly, Multi-agarose plates (MAP) containing an array of 96 agar pads each were produced using acrylic sheets, adhesive sheets and a laser cutter (SI Fig. 2). The assembled plates were filled with solutions of molten agarose in LB with different concentrations of mecillinam, salts or sucrose. The solutions were pre-heated to 80^°^C to let the agarose dissolve and then cooled to 60^°^C before mecillinam was added. The molten agarose was then immediately poured in the wells to minimize mecillinam exposure to high temperature. 1.5 *µ*l of cell suspensions from overnight cultures at a final OD600 between 0.1 and 0.4 were added on top of each pad. The plate was then sealed with a 110× 74× 0.17 mm coverslip, and placed in the focus of a microscope with an enclosure pre-heated at 37^°^C. As before, brightfield imaging was performed using a Nikon 40x CFI Plan Fluor air objective with a numerical aperture of 0.75 and a Teledyne FLIR BFS-U3-70S7M-C camera with a 7.1 MP Sony IMX428 monochrome image sensor.

### Growth rate analysis

To extract maximum growth rates from plate reader curves, we used (24) with the following ranges of fitting parameters: amplitude −5,5; flexibility −6,2; error −5,2 as in (25).

## RESULTS

### Specific salts, not osmolarity, determine survival to mecillinam in *E. coli* Δ*cysB* in LB

Urine with low osmolarity abolishes resistance in Δ*cysB* strains (18). To better understand how high osmolarity supports Δ*cysB* resistance we set out to quantify the effects of various osmolites on the minimum inhibitory concentration (MIC) of BW25113 *E. coli* and its Δ*cysB* derivative in LB medium using the microdilution broth technique in a plate reader. All of the LB based media used in this work are produced starting from a base low salt LB medium which we call LB_*L*_ (*L*, for low salt) which contains 0.5% NaCl. We then add osmolites to the LB_*L*_ taking into account the number of dissociated ionic species such that we maintain the total osmolarity constant among conditions. Further details on concentrations and preparation are given in the *Materials and methods* section and SI Table 1. Using microbroth dilutions in a plate reader, we treated *E. coli* with 13 concentrations of mecillinam at two osmolarities. We repeated the experiment with 14 charged and non-charged osmolites that included salts of monovalent and divalent cations, phosphates, sulphates and sugars (Fig. 1A). We observed a mild protective effect against mecillinam treatment that was *cysB*-independent (Fig. 1B and SI Fig. 3). Δ*cysB*-dependent resistance to high concentrations of mecillinam emerged only in the presence of certain salts, specifically: sodium, potassium and lithium chlorides, and the sulfates of sodium and ammonium. Other chlorine salts with monovalent cations such as ammonium chloride and salts of divalent cations failed to elicit resistance. Resistance was not linked to any specific dissociated ion. Interestingly, neutral species such as sucrose and sorbitol did not induce high level resistance in the Δ*cysB* background in LB (Fig. 1B). In all conditions, osmolites did not significantly alter the morphological defects caused by mecillinam, but influenced cell lysis (Fig. 1C).

**Figure 1.**
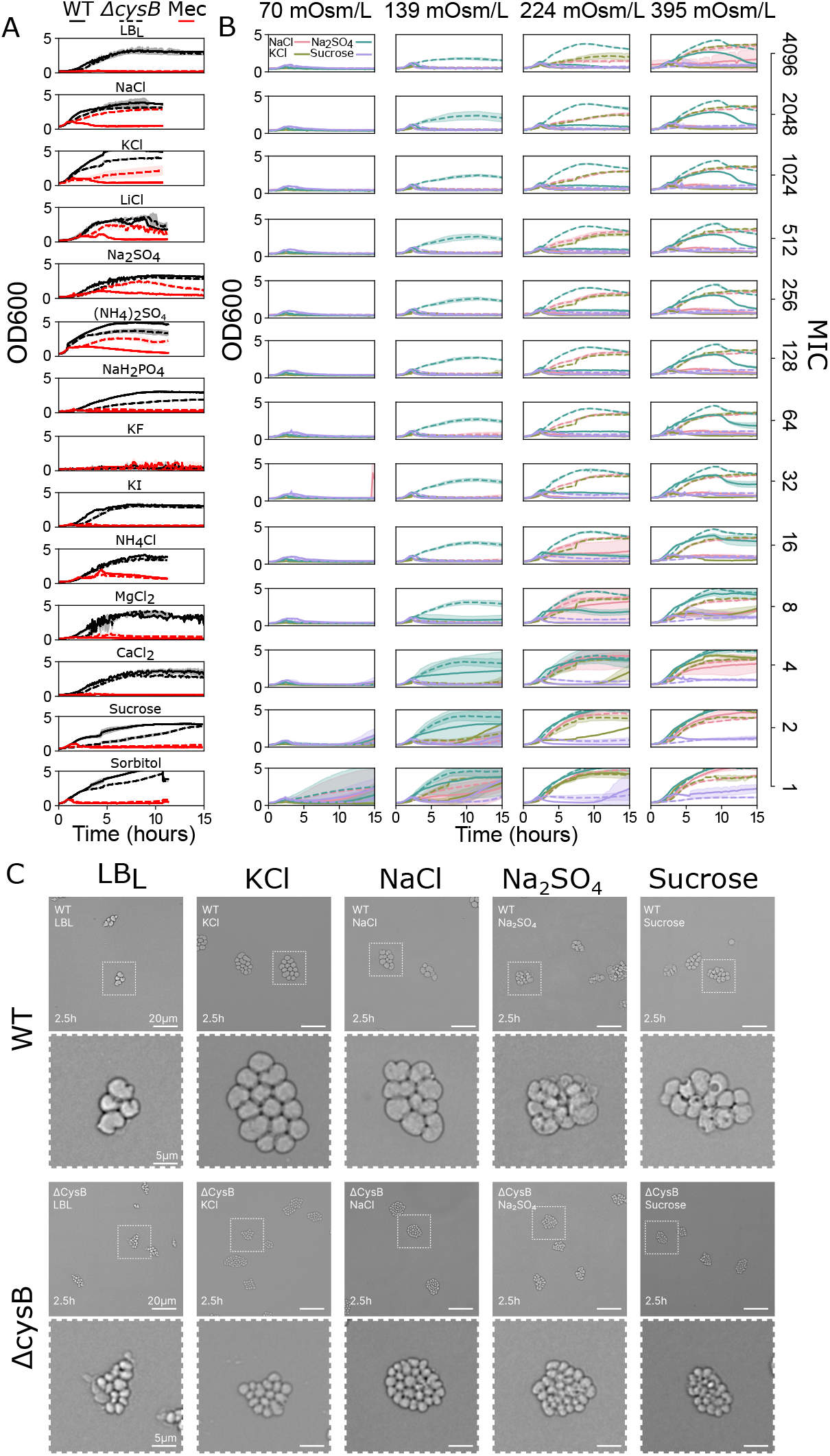
Osmolarity mildly decreases mecillinam effectiveness, while certain salts prompt Δ*cysB* resistance. a) efficacy of 128 *µ*g/ml (red) against Δ*cysB* (dotted line) and reference strain (uninterrupted line) in the presence of different osmolites. LB*_L_* indicates LB with low NaCl (0.5 g/l). NaCl indicates LB*_L_* plus 9.5 g/l of NaCl. All of the other media are produced by adding to LB*_L_* osmolites until the osmolarity of LB + 10 g/l of NaCl is matched. b) MIC changes at different concentrations of mecillinam (bottom to top) and NaCl, KCl, Na_2_SO_4_ and sucrose (left to right). The lines are averages of 3 repeats and the shaded areas indicate the standard deviation. c) microscopy images of cells after 2.5 hours treatment with 1 *µ*g/ml of mecillinam on solid LB*_L_* medium supplemented with the osmotic equivalent of 9.5 g/l NaCl for different osmolites. The second and third rows show zoom-ins of the areas in the first and third rows that are delineated by white dotted rectangles.

### Δ*cysB* also confers salt-independent resistance to novobiocin, but no resistance to other antibiotics

To gain further insights into the salt-dependent mechanism of resistance we tested whether Δ*cysB* cells were resistant to other antibiotics in the same conditions, by comparing the MIC of Δ*cysB* and of the reference strain in LB with low or high salt, across a panel of 7 further antibiotics. We biased our set particularly towards other *β*-lactams, which made up half of our selection, as they are structurally closer to mecillinam. None of the antibiotics tested showed increases in MIC with the exception of novobiocin to which, however, the Δ*cysB* knockout was resistant independently of the salt concentration or medium osmolarity (Table 1, Fig. 2A and B). This suggests that the salt-dependent mechanism of resistance of Δ*cysB* cells is specific to mecillinam, and it is different from the mechanism of resistance to novobiocin.

**Table 1.**
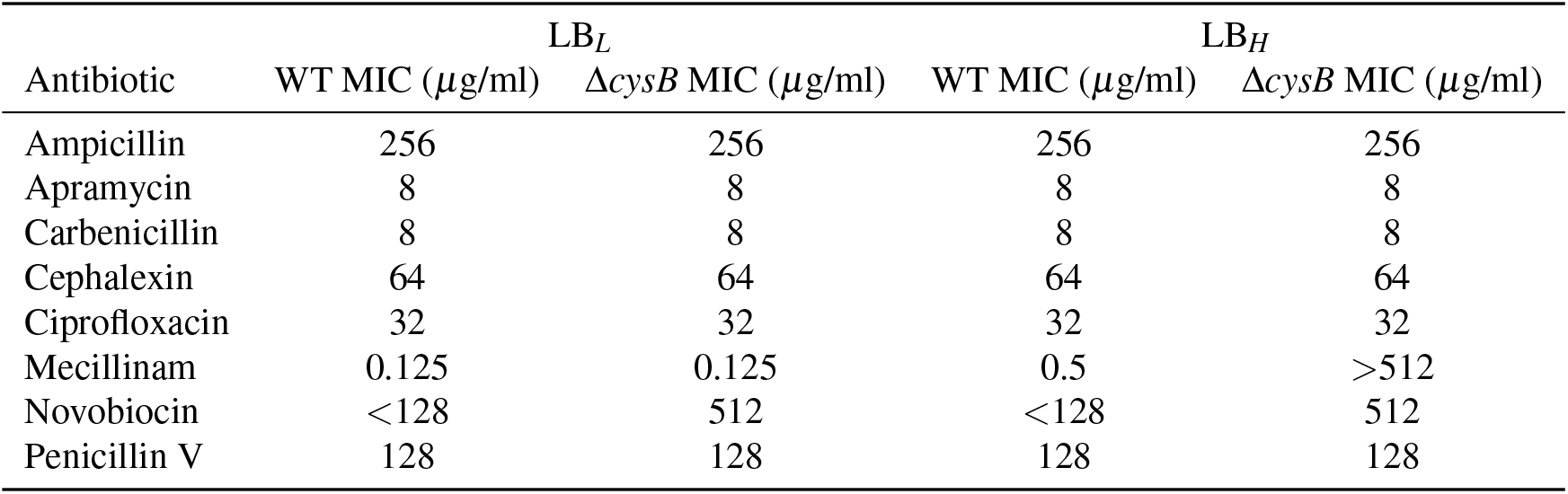
MIC of different antibiotics against reference strain (WT) and *cysB* knockout in LB_*L*_ (0.5 g/l NaCl) and LB*_H_* (10 g/l NaCl).

| Antibiotic | $LB_L$ | | $LB_H$ | |
| --- | --- | --- | --- | --- |
| | WT MIC (μg/ml) | $\Delta cysB$ MIC (μg/ml) | WT MIC (μg/ml) | $\Delta cysB$ MIC (μg/ml) |
| Ampicillin | 256 | 256 | 256 | 256 |
| Apramycin | 8 | 8 | 8 | 8 |
| Carbenicillin | 8 | 8 | 8 | 8 |
| Cephalexin | 64 | 64 | 64 | 64 |
| Ciprofloxacin | 32 | 32 | 32 | 32 |
| Mecillinam | 0.125 | 0.125 | 0.5 | >512 |
| Novobiocin | <128 | 512 | <128 | 512 |
| Penicillin V | 128 | 128 | 128 | 128 |

**Figure 2.**
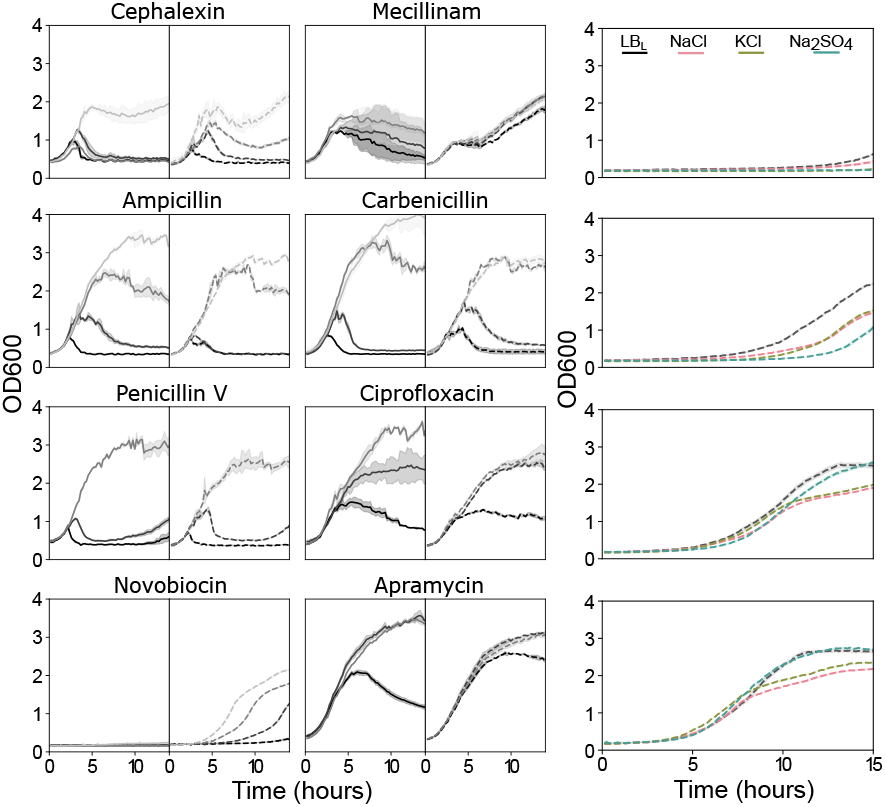
The *cysB* knockout does not confer resistance to other *Β* -lactams and confers it unconditionally to novobiocin. Growth curves from a plate reader showing: a) susceptibility profile of the *cysB* knockout in LB + 10 g/l NaCl against various antibiotics. Black indicates the MIC of the reference strain in LB*H* (Table 1) and different shades of gray indicate sub-MIC concentrations, at progressive 2x dilutions (1:2, 1:4, 1:8). Fainter shades indicate lower antibiotic concentrations. b) Profiles of novobiocin efficacy against the *cysB* knockout in the presence of osmolites that prompt resistance to mecillinam. The concentrations range from 512 *µ*g/ml (top, MIC) to 64 *µ*g/ml (bottom, 1/8 MIC). The lines are averages of 3 repeats and the shaded areas indicate the standard deviation.

### Salt only matters if present during mecillinam treatment, but it does not accelerate mecillinam breakdown

To test whether the protective effect of salt in the Δ*cysB* background could be the consequence of a long-lasting adaptive phenomenon, we evaluated the mutant’s susceptibility to the drug after pre-exposure to high salt medium overnight. When treated in low-salt medium, cells lacking *cysB* were susceptible to mecillinam independently of the pre-culture conditions (Fig. 3A). Salt protection is therefore rapidly lost in the absence of salt. Alternatively, salt could limit mecillinam efficacy by accelerating its degradation rate, although it is unclear how the mutant would participate in degradation dynamics. Indeed, it has recently been shown that the stability of mecillinam can vary across physiologically relevant pH conditions (26). We tested whether salt affects mecillinam stability by incubating the drug in LB_*L*_, LB_*L*_+(10 g/l NaCl), and LB_*L*_+(15.4 g/l Na_2_SO_4_) and adding cells after several 2 hours (Fig. 3B) and 3 hours (Fig. 3C) time intervals as in (26), but we did not find any correlation between inhibitory concentration and antibiotic incubation time over 9 hours of observation (Fig. 3B and C).

**Figure 3.**
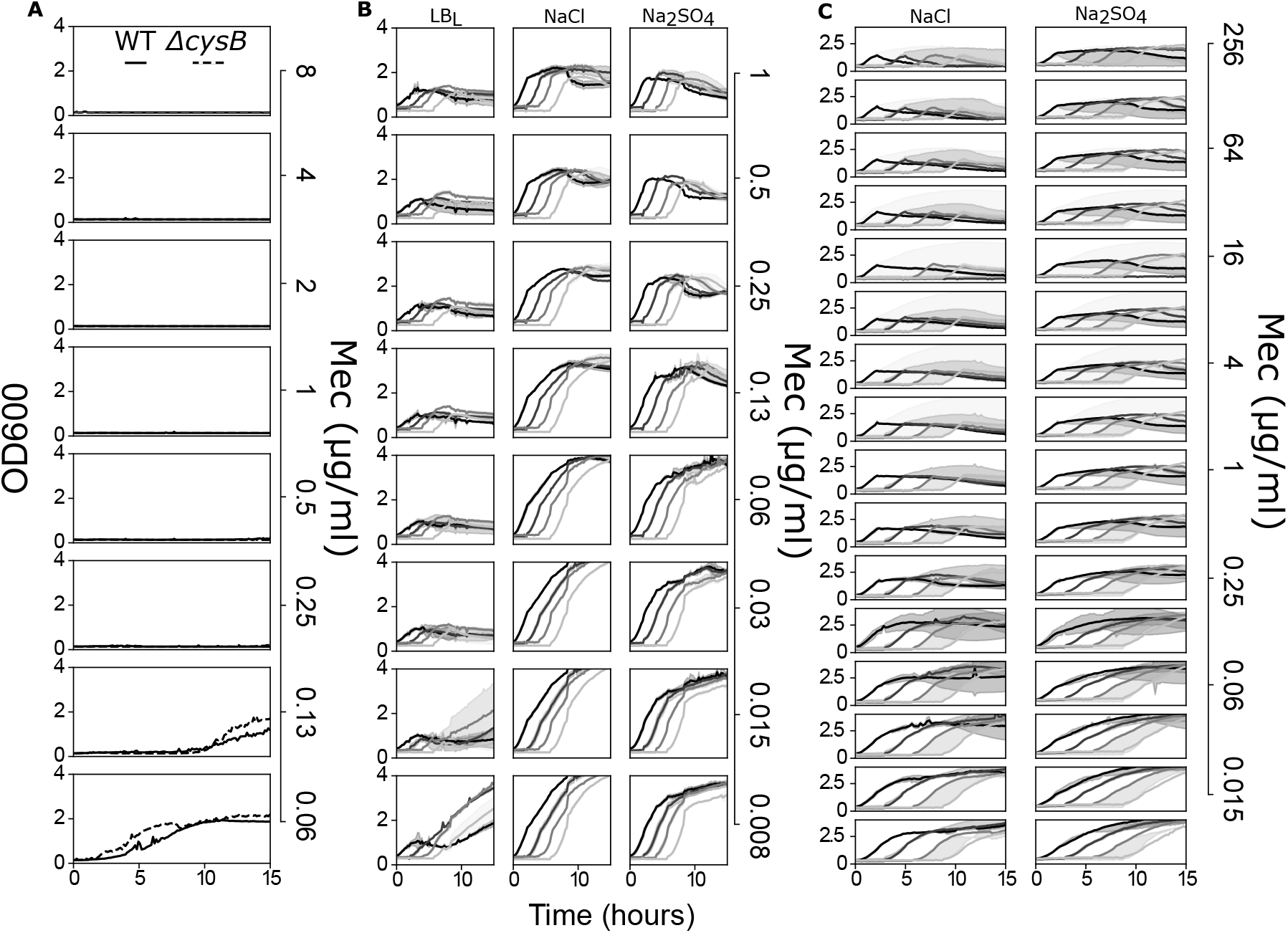
Salt protection has fast dynamics and does not depend on antibiotic breakdown. a) Treatment of reference strain (solid line) and Δ*cysB* (dotted line) with mecillinam in LB*_L_* after overnight growth in LB*H* (10 g/l NaCl). b) Growth curves of the reference strain *E. coli* in LB*_L_*, LB*_L_*+(10 g/l NaCl), and LB*_L_*+(15.4 g/l Na_2_SO_4_) after 0, 2, 4 and 6 hours incubation of mecillinam with the medium in the plate reader at 37°C. Different shades of gray indicate different pre-incubation times (darker indicates 0, lighter indicates 6 hours). c) Growth curves of the reference strain *E. coli* in LB*_L_*+(10 g/l NaCl), and LB*_L_*+(15.4 g/l Na_2_SO_4_) after 0, 3, 6 and 9 hours incubation of mecillinam with the medium in the plate reader at 37 ° C. Different shades of gray indicate different pre-incubation times (darker indicates 0, lighter indicates 9 hours). The lines are averages of 3 repeats and the shaded areas indicate the standard deviation.

### Sodium chloride confers resistance to Δ*cysB E. coli* in urine

We then sought to characterize the impact of salt on the conditional Δ*cysB* resistance to mecillinam in urine. In pooled, filtered human urine, *E. coli* did not show robust growth and, as expected based on our previous observations on slow growth media (16), mecillinam was inefficacious (Fig. 4A). Susceptibility emerged upon supplementation of components of defined media (Fig. 4B) to urine (Fig. 4C and D), with a sharp transition between growth rates 0.5 and 0.6 h^*−*1^ (Fig. 4D). While EZ (a commercial mix of amino acids and vitamins (Teknova, US)) was the main driver of growth rate increases, such transition was independent from any specific medium components which were found on both sides of the threshold growth rate. In defined lab medium, NaCl had no effect on the MIC of the reference strain, while it prompted Δ*cysB*-dependent resistance as we had observed in LB (Fig. 4B). Interestingly, in urine enriched with NaCl, the Δ*cysB* strain still showed resistance to mecillinam compared to the reference strain, but this was due to increased susceptibility of the latter (Fig. 4C). Na_2_SO_4_ did not provide protection in urine (SI Fig. 4). Although this did not change the overall trends, we also noted that the use of yeast extracts from different brands during pre-culture in LB had an effect on growth in urine (SI Fig. 4).

**Figure 4.**
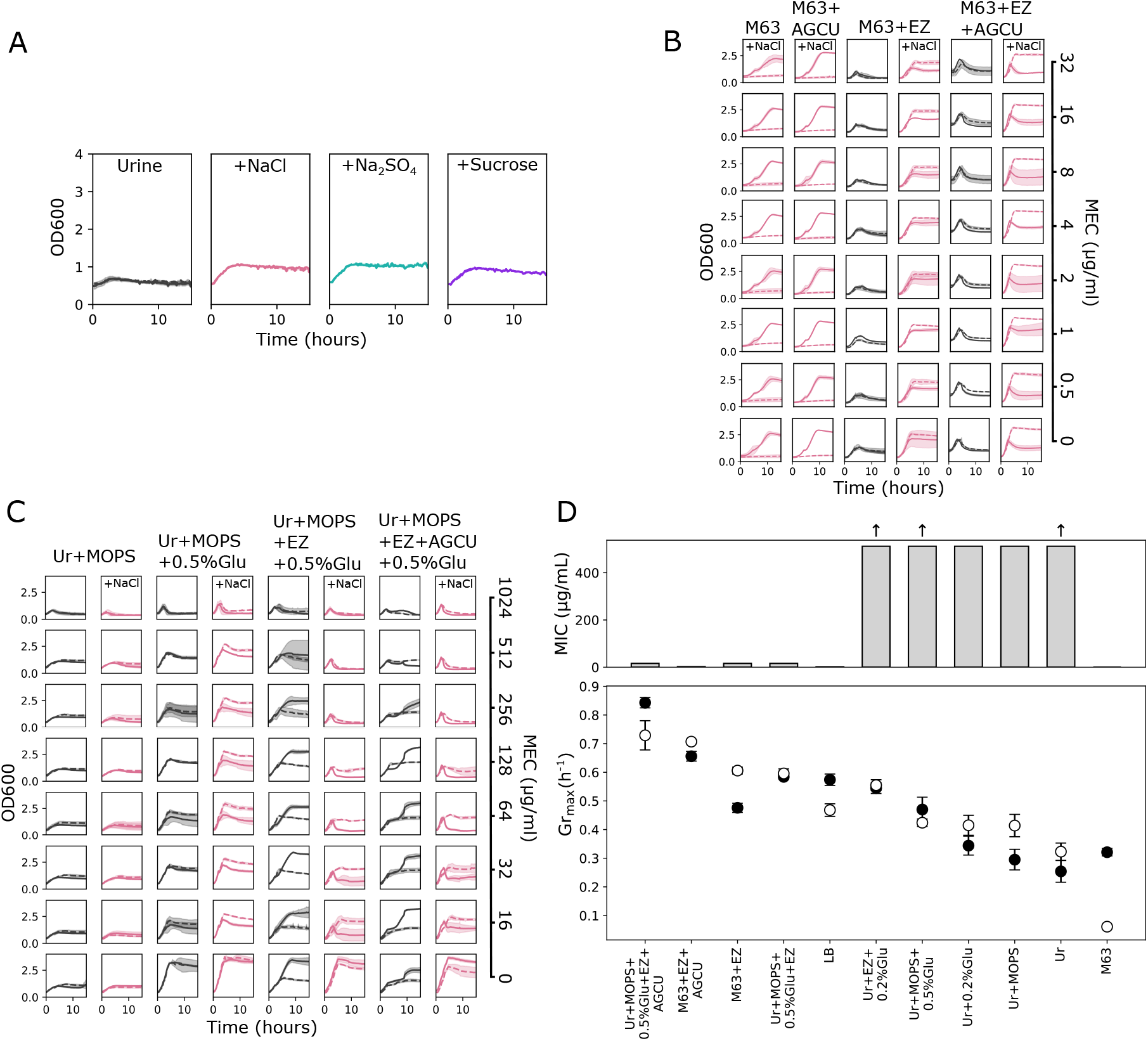
Mecillinam is scarcely efficacious in urine at low growth rates and salt-dependent resistance emerges when nutrients are added. a) Growth curves of reference strain (uninterrupted line) and *cysB* knockout (dotted line) in the presence of various osmolites (obtained as LB*_L_* plus the osmotic equivalent of 9.5 g/l) and 512 *µ*g/ml mecillinam. b-c) Growth curves of reference strain (uninterrupted line) and *cysB* knockout (dotted line) in the presence of different mecillinam concentrations. Curves in pink show experiments with 9.5 g/l NaCl added to LB*_L_*, LB*_L_* is in black. Different media: M63, M63+AGCU, M63+EZ and M63+EZ+AGCU (b), and urine+MOPS, urine+MOPS+0.5% glucose, urine+MOPS+EZ+0.5% glucose, and urine+MOPS+EZ+AGCU+0.5% glucose (c), are indicated at the top. d) MIC of the referemce strain (top) and maximum growth rates (bottom) for reference strain (black circles) and *cysB* knockout (white circles). The arrows indicate conditions in which the MIC might be higher (pointing up), but was not measured further. For all media, MICs and growth rates were extracted in the presence of 10 g/l of NaCl. Growth conditions are ordered in descending order based on the highest maximum growth rate of the faster growing strain. Growth curves are averages of 3 repeats and the shaded areas indicate the standard deviation. Maximum growth rates are obtained from at least 3 replicates and the error bars indicate the standard deviation.

### Δ*cysB* cells are smaller and grow slower than the wild-type on solid LB medium

Cell size is among the properties that can change within the timescale of a single cell generation, therefore fitting with the time dynamics demonstrated by the salt-dependent mechanism and we have recently demonstrated that small cells are less susceptible to mecillinam (16). A salt-induced reduction of the cell size below the bactericidal threshold could explain conditional survival. Therefore, we sought to compare the morphological parameters of the Δ*cysB* strain with those of the reference strain across different concentrations of NaCl, KCl, Na_2_SO_4_ and of sucrose using microscopy. To image cells with high-throughput across conditions we used our recently developed multi-agarose pad (MAP) (SI Fig. 2) (22; 23). In all of the conditions tested, Δ*cysB* cells were smaller than those of the reference strain (Fig. 5, SI Fig. 5B). In addition, for both strains, the addition of osmolites reduced cell size causing shortening and mild widening (Fig. 5M). Because reduced cell size is associated with mecillinam survival, we hypothesize that this is an additional mechanism, further to cytoplasmic blebs stabilization, through which increased medium osmolarity protects cells from mecillinam as observed in Fig. 1B and SI Fig. 3. However, this is not the mechanism by which Δ*cysB* strains attain their salt-dependent resistance because both sucrose and salts cause size reduction (Fig. 5A-J). On solid LB medium, Δ*cysB* cells grew slower than the reference ones (Fig. 5N and O,, SI Fig. 5A) in agreement with the growth rate measurements performed in liquid for the same medium (Fig. 4D).

**Figure 5.**
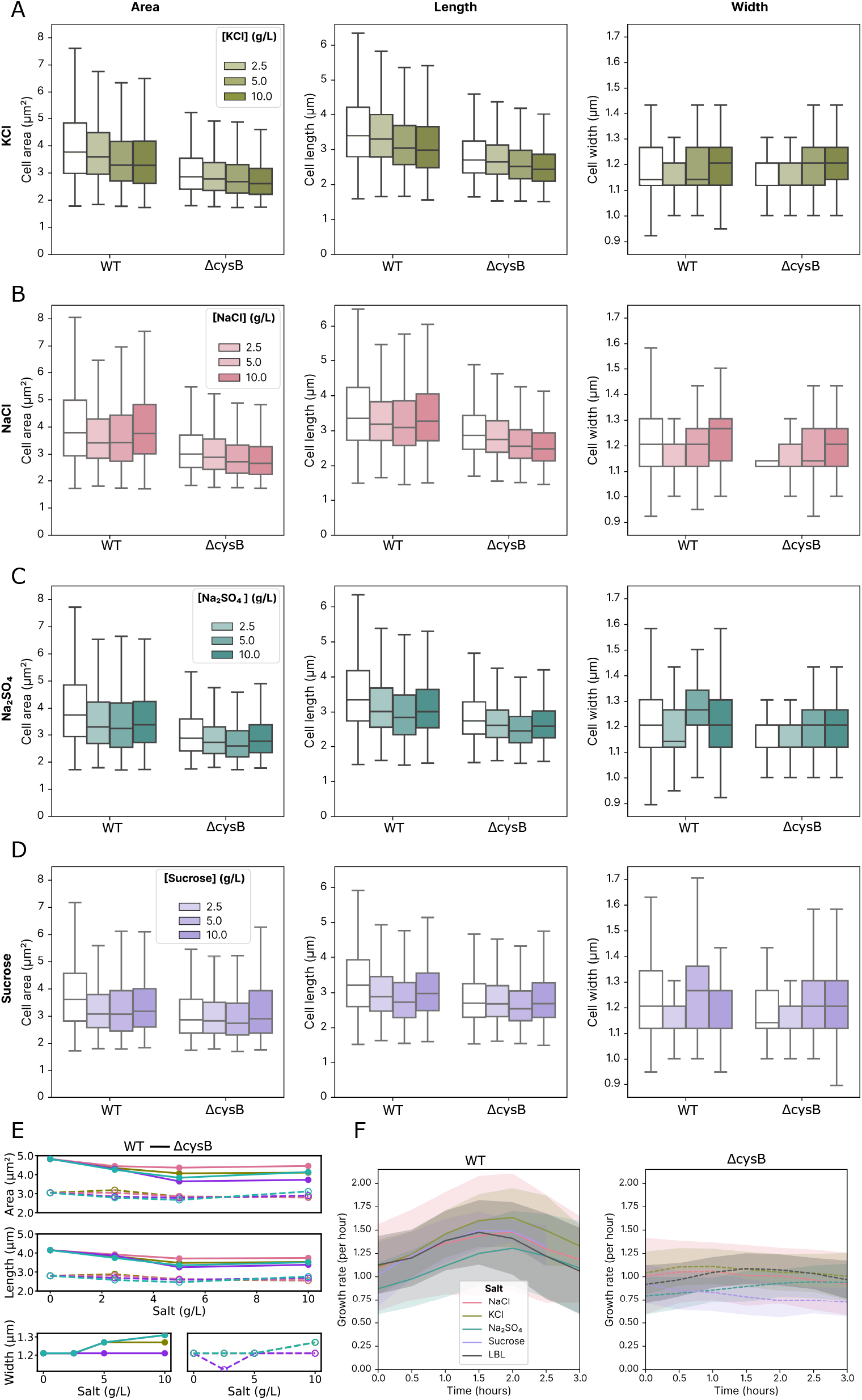
Δ*cysB* grows slower than reference strain on solid LB medium and salts decrease the cell size of both. a-d) single cell measurements on reference and Δ*cysB E. coli* in media supplemented with KCL (a), NaCl (b), Na_2_SO_4_ (c) and sucrose (d). Single cells are collected from at least 3 replicate experiments. Bar plots indicate median, first and third quartile, as well as the standard deviation.

## DISCUSSION

Using microbroth dilution and microscopy, we have shown that the Δ*cysB* mediated conditional resistance of *E. coli* to mecillinam is salt-dependent rather than osmolarity-dependent. Furthermore, not all salts enable resistance. Among the salts tested, sulfates and several but not all chlorides of monovalent cations induced resistance. We did not observe resistance for phosphates nor for chlorides of divalent cations. Resistance did not depend on the presence of any specific ion. We have shown that the mechanism of salt-dependent resistance is specific to mecillinam, it does not affect sensitivity to other *β*-lactams, and it is different from the resistance mediated by Δ*cysB* to novobiocin. Further, we have shown that the impact of salt is transitory, and resistance depends on the simultaneous presence of mecillinam and salt. However, as also indicated by the fact that salt does not robustly protect the reference strain, we have observed no acceleration in mecillinam breakdown due to salt. We have then demonstrated that the protective effect of salt is not limited to LB medium, but it extends to other commonly used defined media and to human urine as long as these are able to support a sufficiently high growth rate and does not depend on a single nutritional factor. Lastly, by measuring cell morphological parameters across the various conditions, we have shown that growth at high osmolarity decreases cell size.

Contextualizing these results to our recent discovery that slow growing, small cells survive mecillinam treatment, we are able to add further nuance to previous statements suggesting that increased fluid intake should be recommended during mecillinam treatment. Indeed, while fluid intake will decrease osmolarity and salt content in urine, it will also decrease the concentration of nutrients, potentially shifting infections towards growth regimes that require very high concentrations of mecillinam to be effective. Indeed, our Fig. 4C shows that reference lab strains can grow at concentrations of mecillinam that surpass 0.5 mg/ml in pooled human urine, a level that is rarely achieved and only briefly maintained in patients (27). Reducing sodium consumption seems therefore the preferable strategy.

While the mechanism of resistance of Δ*cysB E. coli* remains to be determined, our work expands the characterization of the knockout’s phenotype, suggesting the involvement of cellular ionic balance. Ionic fluxes are central to osmotic control (28) and the role played by cysteine in the oxido-reductive balance of the periplasmic space (29; 30) might be linked to these fluxes and the behaviours of transporters. Osmotic balance is, in turn, well known for its involvement in *β*-lactam bactericidal activity (9; 11). Additionally and not necessarily alternatively, salts and other osmolites influence protein conformation and interactions (31), including those of FtsEX (32) which makes up *E. coli*’s divisome, a complex that is central to cell wall homeostasis. An altered ion homeostasis might impinge on the behavior of the divisome, already affected by the morphological changes induced by mecillinam, leading to unexpected phenotypes.

## AUTHOR CONTRIBUTIONS

L.M. conceived the experiments. L.M. and M. K. performed the experiments and analyzed experimental data. L.M. interpreted the results and wrote the original draft. J.K. supported the work on imaging. L.M., M.K., J.K. and P.C. wrote the final manuscript.

## ACKNOWLEDGEMENTS

We thank Abimbola Feyisara Adedeji-Olulana, Jamie Hobbs, Marco Mauri, Rosalind Allen, Jacob Biboy and Waldemar Vollmer for helpful discussions. This work was financially supported by the Herchel Smith postdoctoral fellowship to L.M., the EU EC ITN Phymot, Marie Sklodowska-Curie grant agreement No 955910 to M.K. and P.C. and the UKRI EP/T002778/1 to L.M., J.K. and P.C..

## COMPETING INTEREST

None declared.

## DATA AND MATERIALS AVAILABILITY STATEMENT

All data is deposited at 10.5281/zenodo.15652483.

